# Multicolor single particle reconstruction of protein complexes

**DOI:** 10.1101/255364

**Authors:** Christian Sieben, Niccolò Banterle, Kyle M. Douglass, Pierre Gönczy, Suliana Manley

## Abstract

Single-particle reconstruction (SPR) from electron microscopy images is widely used in structural biology, but lacks direct information on protein identity. To address this limitation, we developed a computational and analytical framework that reconstructs and co-aligns multiple proteins from 2D super-resolution fluorescence images. We demonstrate our method by generating multi-color 3D reconstructions of several proteins within the human centriole and procentriole, revealing their relative locations, dimensions and orientations.

---

Macromolecular complexes within cells usually contain multiple protein species, whose precise arrangement is required to generate properly functioning molecular machines. Single particle analysis of electron microscopy (EM) images has been used to build 3D reconstructions of such complexes, recently with near-atomic resolution^1,2^. To deduce the spatial organization of specific proteins, computational methods have been used to dock structures deciphered from X-ray crystallography or NMR within 3D reconstructed particles^1,3^. Alternatively, immunogold or nanobody labelling can reveal the location and conformational state of target proteins^4,5^, whereas electron density map differences can provide information on the position of mutated or missing proteins^6^. Nevertheless, it remains challenging to locate native proteins within 3D reconstructions, which is essential for deciphering the architecture underlying the assembly mechanisms and functional modules of macromolecular complexes.

Fluorescence-based single-molecule localization microscopy (SMLM) can help to address this challenge, as demonstrated for the nuclear pore complex using 2D averaging^7^. Extending to 3D, a recent implementation of single-particle reconstruction (SPR) from 2D SMLM images demonstrated isotropic reconstruction on DNA origami and simulated data^8^. However, multi-color particle reconstruction of actual macromolecular complexes requires generating large image libraries of multiple proteins and solving the complex problem of 3D multi-channel alignment. Here, we developed a systematic framework that addresses both of these challenges. We used a dedicated high-throughput SMLM setup^9^ to generate the large multi-color particle data sets required for SPR, which we then processed using a semi-automated computational workflow to reconstruct and align multiple proteins onto a single 3D particle. We applied our method to the human centriole, and adapted it to accommodate off-axis structures, such as those important during procentriole formation. Centrioles are evolutionarily conserved sub-diffraction limited cylindrical organelles that seed the formation of cilia, flagella and centrosomes^10^. The mature human centriole comprises nine-fold symmetrically arranged microtubule triplets and contains >100 different proteins organized into distinct substructures^11^. For instance, distal appendages harbor the protein Cep164 and are key for cilium and flagellum formation. Moreover, a torus encircling the proximal part of the mature centriole and comprising the proteins Cep57/Cep63/Cep152 acts as a nucleation site for the emerging procentriole, whose assembly relies on the self-organization of the HsSAS-6 protein into a cartwheel^12,13^. The exact dimensions of components within the Cep57/Cep63/Cep152 torus and the position of the procentriole with respect to this torus remain unclear.

To demonstrate multicolor 3D SMLM reconstruction, we imaged protein species within centrioles and procentrioles (see experimental workflow in Supplementary Fig. 1). Centrosomes were isolated from human KE37 cells arrested in S phase, concentrated onto coverslips by centrifugation, followed by immunolabeling and dual-color fluorescent staining (Supplementary Note 1). Thereafter, we used high-throughput SMLM^9^ to image on average 150 centrioles per field of view (Supplementary Fig. 2). Localizations belonging to centrioles were segmented using a mask generated through automated OTSU thresholding of the widefield images. A density-based filter (DBSCAN^14^) was then applied to separate adjacent centrioles (Supplementary Fig. 3). Only densely labelled centrioles (typically 10-20% of the initial dataset) were rendered and used to populate the particle dataset. Further particle processing was performed as follows using EM routines integrated into Scipion^15^ (Supplementary Note 2). Dual-color particles (Supplementary Fig. 4) were classified and class-averaged using template-free maximum-likelihood multi-reference refinement (ML2D)^16^. Due to the high degree of radial symmetry within centrioles^10^, a low number of classes (typically 8-15) were chosen, thereby reducing computational complexity. Nevertheless, this approach is also applicable to particles of unknown symmetry, as verified *in silico* (Supplementary Fig. 5). The class averages best resembling the input particles (Supplementary Note 2) were then used to compute an initial 3D model followed by structural refinement based on matching its 2D projections to the input particles. In this manner, we reconstructed the torus protein Cep152, which we found to exhibit a ~260 nm diameter (Fig. 1a), consistent with the ~242 nm value measured for SNAP-Cep152 by STED microscopy^17^. In addition, our 3D reconstruction enabled us to determine for the first time that the height of the torus is ~190 nm. Following the same procedure, we reconstructed the well-known bacteriophage T4 (Supplementary Fig. 6), thus demonstrating the flexibility of this 3D SMLM reconstruction pipeline.

**Figure 1:**
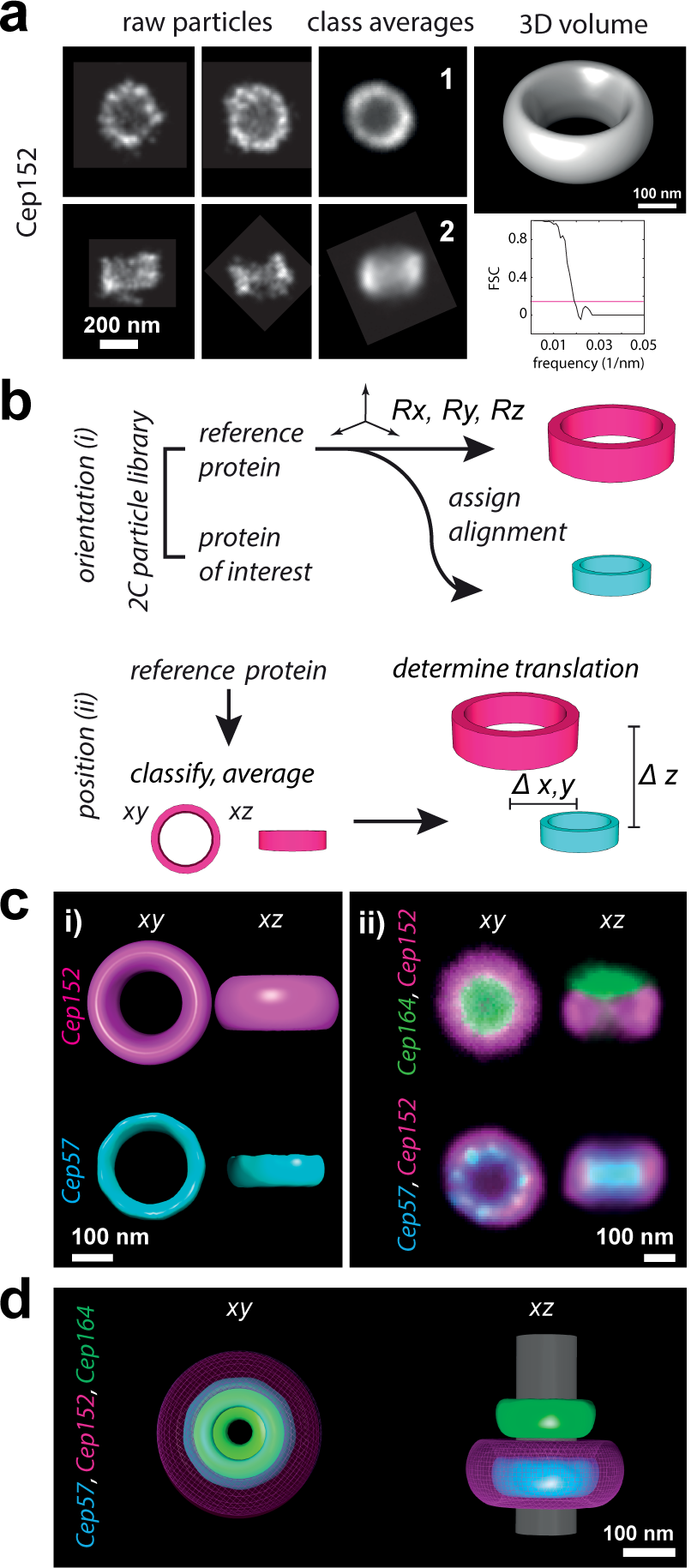
Multi-color single particle reconstruction. (**a**) Purified human centrosomes were immunolabelled against Cep152 and Cep164 and imaged using high-throughput SMLM to collect 2306 single dual-color particles. Shown are examples of input raw particles (Cep152 channel) and the corresponding class averages (1, 2). A 3D volume model (top right) was reconstructed at a resolution of 52 nm as assessed by Fourier shell correlation (FSC) (lower right). (**b**) The workflow for multi-volume alignment from two-color (2C) SMLM particles is divided into two steps: (i) orientation and (ii) position alignment. (**i**) The two imaged proteins belong to the same structure and are thus in the same orientation, so we identified the orientation of only one (reference protein, magenta) and assigned it to the protein of interest (cyan). (**ii**) To correctly position the two resulting co-oriented volumes with respect to each other, reference protein particles were identified in side view. By collapsing one axis (x or y), we could determine the axial translation ( z) between the two imaged proteins owing to the symmetry of the centriole. **(c)** (i) Orientational alignment of the two centriolar proteins Cep57 (cyan)/Cep152 (magenta) results in two co-oriented volumes shown in top (xy) and side (xz) views. (ii) Translational alignment of Cep152 (magenta)/Cep164 (green) and Cep152 (magenta)/Cep57 (cyan) results in two 2-color reconstructions. **(d)** The resulting three-color volumetric map is shown in bottom (xy) and side (xz) views, with a volume of the known dimensions of the microtubule outer wall (grey) as reference.

To achieve multi-color reconstruction, we first considered the case of proteins sharing a principal symmetry axis. We collected dual-color images of Cep152/Cep164, as well as of Cep152/Cep57, keeping Cep152 as a reference to combine both datasets into a single 3D map. This reduced the problem of alignment to only two 3D volumes at once. We divided the alignment process into two steps: i) co-orient both particles and reconstruct their volumes; ii) co-align the volumes along the symmetry axis (Fig. 1b). Since both proteins are integrated into the same structure, the corresponding particles share the same orientation. Therefore, it is sufficient to find the orientation of particles in one channel (i.e. the reference), then conserve and assign the alignment parameters to the second channel. Given the challenge of imaging two proteins with high resolution, which is typically limited by protein abundance and/or labeling efficiency, this procedure offers the great advantage that only the imaged reference protein needs to contain enough information to be oriented. Following this procedure results in two co-oriented volumes.

To co-align protein volumes on a shared symmetry axis, one only needs to align their top (xy) and side (xz) view projections (Fig. 1b, Supplementary Fig. 7). We streamlined this process by using supervised machine learning (MATLAB, Classification Learner) to identify top and side view projections from a combination of 12 calculated shape descriptors (Supplementary Note 3). The models were trained on ~10% of particles and successfully identified ~85 % of side view projections. This method offers the advantage that after having been trained for one reference protein (e.g. Cep152), the model can readily be applied to other datasets exploiting the same reference, greatly facilitating axial alignment. We typically extracted 50–100 side view particles per imaged protein pair. To overlay the particles, we implemented an additional 2D averaging step comprising particle rotation in 1° increments followed by translational alignment and cross-correlation to find optimal overlap between particle pairs. Again, the 2D alignment parameters for the reference protein can be applied to the other imaged protein. This approach automates the computation of Δz while enabling a more precise estimate of particle dimensions (Supplementary Fig. 8).

Importantly, this streamlined workflow allowed us to reconstruct and co-align Cep57, Cep152 and Cep164 in a three-color volumetric map of the human mature centriole (Fig. 1d). This reconstruction revealed that whereas the Cep57 torus is aligned axially with that formed by Cep152, as expected from their known association in cells^17^, it has smaller dimensions (~230 nm in diameter and ~130 nm in height), placing it close to the outer microtubule wall. Interestingly, we also discovered a previously unidentified nine-fold radially symmetric distribution of Cep57 (Supplementary Fig. 8), further suggesting association with the nine-fold symmetrical outer microtubule wall of the centriole, perhaps via its microtubule binding domain^18^. We also confirmed Cep164’s previously observed nine-fold symmetric organization, while locating its N-terminus more proximally and closer to the centriolar wall than previously reported^19^ (see also Supplementary Fig. 9).

The above approach works well for proteins positioned on the same principal symmetry axis, but there are important cases for which this does not hold. To exemplify this point, we extended our method to the procentriole, marked by the protein HsSAS-6, which emerges from a single focus on the torus containing Cep57/Cep63/Cep152 during organelle assembly^20^. We collected dual-color images of Cep152/HsSAS-6, and generated average top and side views following the procedure described above (Supplementary Fig. 7). In this case, the orientation of Cep152 was insufficient to define that of HsSAS-6, since the two proteins do not share a symmetry axis (Fig. 2a). One solution would be to combine the images from both proteins into a single channel and perform class averaging and alignment on the resulting dataset. However, the signal from the larger Cep152 structure would dominate and prevent alignment of the smaller HsSAS-6 volume. Instead, we combined the information from the two channels by fusing both rendered images in a weighted sum, where the HsSAS-6 signal was given twice the weight of Cep152. We then used the fused particles for structural refinement of the initial Cep152 volume without any symmetry constraint (Fig. 2b), and fit the individually reconstructed protein volumes into the asymmetric global structure to achieve two-color volumetric reconstruction of the growing procentriole in the context of its centriole (Fig. 2c). Interestingly, we found that the combined reconstruction displays a lower resolution than the individually resolved structures (Fig. 2c), indicative of a flexible relative positioning of the two entities. Indeed, we found the angle θ between the two measured from individual side views (Fig. 2a) to be variable, with an average value of 15.5± 2.7 (SEM, n = 25), in agreement with our 3D reconstruction. Together, these findings indicate a loosely defined orientation between the torus and the emerging procentriole, consistent with suggestions from some EM analysis^21^.

In conclusion, we developed a novel framework that generates multi-component 3D volumes from dual-color 2D SMSM datasets, and demonstrated it to construct a three-component 3D model of the human centriole and procentriole, thus revealing novel features of their architecture. This method is a flexible workflow that can easily be adapted to other multiprotein complexes and imaging modalities. Combining information from 3D SMLM reconstructions with EM particle reconstructions will likely prove invaluable in the future and so will improvements in labelling to permit higher fidelity of fluorescence to the underlying protein structure.

**Figure 2:**
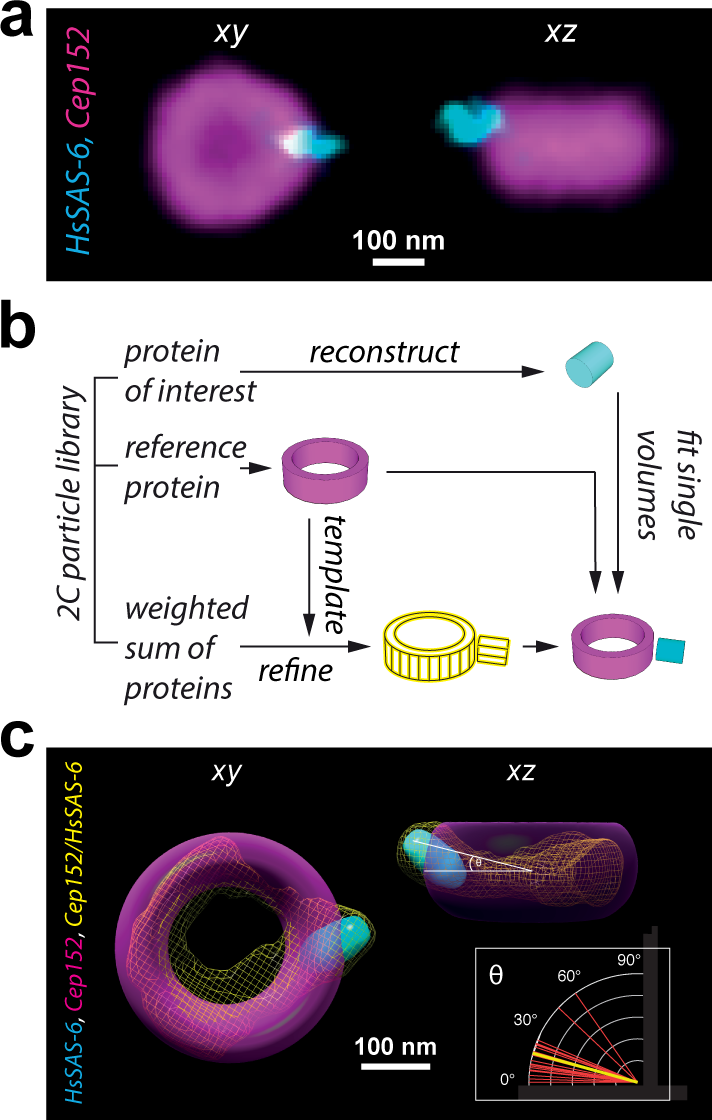
Multi-color single particle reconstruction of an asymmetric protein complex. (**a**) Top (xy) and side (xz) view averages for HsSAS-6 (cyan) and Cep152 (magenta) display a protruding, off-axis HsSAS-6 density. (**b**) Overview of procedure for three-dimensional reconstruction of an asymmetric assembly. First, both the reference protein (magenta) and the protein of interest (cyan) are reconstructed individually, then the joint density (yellow mesh) is reconstructed using the weighted sum of the individual channels with the reference volume as initial template. The individual volumes are then fit into the joint density map to obtain the final asymmetric two-color reconstruction. (**c**) Reconstruction of Cep152 and HsSAS-6 assemblies. The individual volumes of Cep152 (magenta) and HsSAS-6 (cyan) were fit into their joint density map (yellow mesh). The final 3D arrangement shows a non-orthogonal orientation of HsSAS-6 with respect to the toroidal surface of Cep152. Inset: the average orientation of HsSAS-6 arises from a broad distribution of angles measured from 2D aligned side views (red, individual particles; yellow, average angle (θ = 15.5 ± 2.7°)).

## Methods

### Materials and Sample Preparation

Samples were imaged on gold-embedded fiducial cover slips (custom 18 mm, Hestzig). Imaging buffer components were purchased from Sigma. Additional gold fiducials were obtained from Corpuscular (C-Au-0.1) and diluted (1:5) in 0.1 % poly-L-lysine (Sigma) before application. To create a bead sample for two-channel registration, glass cover slips (1.5, Menzel, 25 mm) were plasma cleaned, coated with 0.1 % poly-L-lysine (Sigma) for 30 min and incubated with FluoroSpheres (Dark Red, F8789, Life Technologies) diluted (1:50,000) in water for 10 min. Human centrioles were purified from KE37 incubated for 24h with thymidine following a standard protocol ^22,23^ and spun (10 min at 10,000 g in Corex tubes, JS13.1 Beckman swinging rotor) in 10 mM K-Pipes on gold-embedded fiducial cover slips (custom 18 mm, Hestzig) using a custom centrifuge concentrator, followed by methanol fixation (5 min at −20°C). Samples were then immunostained by overnight incubation at 4° C with primary antibodies (Supplementary Table 1), diluted 1:500 (in PBS supplemented with 1% BSA and 0.1% Tween 20), then washed three times 15 min in PBS and incubated with secondary antibodies coupled with Alexa 647 or DyLight 755 for 1h at room temperature. Finally, the samples were washed again three times for 15 min and stored in the dark at 4 °C until further use.

Bacteriophage T4 was grown and purified following established procedures^24^. To characterize the purified sample, phages were spotted on mica and imaged using atomic force microscopy (JPK Nanowizard). To achieve all-protein labelling, phages were incubated with Alexa 647 NHS-Ester (Life Technologies) (final concentration 10 µM in phosphate buffer, pH 8) overnight at 4° C. The labelled phages were separated from unbound dye using a NAP-5 (GE Healthcare) size exclusion column and stored in the dark at 4 °C until further use. Before SMLM imaging, phages were adsorbed on plasma-cleaned glass cover slips (1.5, Menzel, 25 mm) after coating with 0.1 % poly-L-lysine (Sigma) for 30 min.

### High-throughput SMLM

Two-color SMLM imaging was performed using a flat-field epi illumination microscope^9^. Briefly, two lasers with wavelengths of 642 nm (2RU-VFL-P-2000-642-B1R, MPB Communications) and 750 nm (2RU-VFL-P-500-750-B1R, MPB Communications) were used to switch off fluorophores in the sample, while a 405 nm laser (OBIS, Coherent) controlled the return rate of the fluorophores to the fluorescence-emitting state. A custom dichroic (ZT405/561/642/750/850rpc, Chroma) reflected the laser light and transmitted fluorescence emission before and after passing through the objective (CFI60 PlanApo Lambda Å~60/NA 1.4, Nikon). After passing the respective emission filter (ET700/75M, Chroma or ET810/90m, Chroma), emitted light from the sample was imaged onto an sCMOS camera (Prime, Photometrics). The sample was excited with laser output power of 1200 mW (642 nm) and 500 mW (750 nm). The 405 nm laser was operated with laser output power 1-10 mW. Axial sample position was controlled using the pgFocus open hardware autofocus module (http://big.umassmed.edu/wiki/index.php/PgFocus). Typically, 30-60k frames at 10 ms exposure time were recorded for each field of view using Micromanager^25^. Single- and dual-color SMLM imaging was performed using an optimized SMLM buffer as described previously^26^. See Supplementary Note 1 for more details on the choice of the fluorophores and buffer generation.

### Single-Fluorophore Localization, Channel registration, Drift Correction

Image stacks were analyzed using a custom CMOS-adapted analysis routine (adapted from^27^). Alignment of the Alexa 647 and DyLight 755 data sets was carried out in three steps. We first calculated an affine transformation function from images of fluorescent beads (see **Materials and Sample preparation** section above) acquired in both channels. This first step corrects for differences and aberrations (rotation, magnification) of the emission path between both detection channels. During the next step, both datasets were independently drift-corrected using gold fiducials visible in both channels. For each field of view, we selected 3–6 fiducial markers and used their average trajectory for drift correction. We further extracted bead trajectories from both channels to use them for the third correction step, during which the centroids of the fiducial markers in both channels were matched, resulting in a final lateral translation. Steps one and three were only calculated for and applied to the DyLight 755 data set. Our channel correction procedure resulted in a residual error of on average 25 nm per field of view of typically 80 x 80 µm. Gold fiducial-based drift correction was performed using the open source data management and analysis environment B-Store^28^. All other processing steps were performed in MATLAB 2016a (Mathworks) and are available as part of the supplementary software package.

### Particle Extraction and 3D Reconstruction

Following channel registration, the two localization data sets were ready for particle extraction. The localization maps for each field of view were loaded into MATLAB together with the corresponding wide-field images taken prior to the SMLM stack acquisition. One of the two widefield (WF) images was used for automatic OTSU segmentation to identify the location of individual particles within each field of view. Each identified segment was expanded into a slightly larger rectangle to account for any mismatch between WF image and localization data. The size of the bounding rectangle was chosen dependent on the radius of the segmented region. For diameter < 3 pixel, we expanded the region by a factor of 3, for diameter between 3 and 5 pixel by a factor of 1.5, and for diameter > 5 pixel, we did not expand the region any further. The localizations from both channels were extracted for each segmented particle. Particles were filtered for a minimum number of localizations (typically >100) to ensure good dual particle labelling. At this point, we also applied an upper cut-off to reject too large particle clusters. During the next step, adjacent particles within the same segment were separated using density-based clustering (DBSCAN^14^). An example is shown in Supplementary Fig. 3. We then calculated a number of particle quality and shape descriptors (Supplementary Note 3), as well as the resolution (using Fourier ring correlation^29^) for each particle, which allowed for efficient particle filtering and classification. Finally, particles from both channels were rendered into a pixel image using a 2D histogram function with a bin size of 10 nm and blurred using a Gaussian filter with σ corresponding to the measured localization precision. The final image approximates the probability density distribution of the fluorescent labels on the underlying structure and is a widely-used approach to visualize SMLM data. The particle images were stitched together using the Montage function in ImageJ (Miji for MATLAB) resulting in the final input image for the 3D reconstruction (example shown in Supplementary Fig. 4).

Single particle reconstruction was performed using Scipion, a freely available software package that integrates several widely-distributed and well-developed 3D EM particle reconstruction routines^15^. A brief tutorial of the required steps is provided in Supplementary Note 2. The particle montage images were first imported into Scipion. Depending on the size of the dataset, each montage contained only ~500 particles, which consequently required the generation and import of multiple montages. Since the particles were well aligned, they could be picked and extracted automatically. The particles were aligned again using CL2D (Xmipp3) as the picked region was usually not exactly centered over each particle. The particles were classified using template-free multi reference maximum likelihood (ML2D, Xmipp3) or 2D clustering (CL2D, Xmipp3) classification. Class averages that resembled well the input particles were used (see Supplementary Note 2) to generate the initial model. For symmetric centriolar reconstructions (Cep164, Cep57, Cep152), we selected between 8-12 classes, applying rotational or nine-fold symmetry (Supplementary Note 2). Likewise, for bacteriophage T4, we selected 10 class averages and calculated the initial model using rotational six-fold symmetry (c6). Initial models were calculated using Xmipp2 or Eman2 providing similar results. Finally, the best initial model was refined using particle back projection (Xmipp3). Fourier shell correlation (FSC) was calculated as an output function within the last refinement step (particle back projection, Xmipp3). For the reconstruction of two proteins, we first reconstructed the reference protein using the steps described above, then apply the final alignment to the extracted particles of the protein of interest (function alignment assign), from where the final 3D model can be generated. Please see Supplementary Note 2 for a detailed description. The asymmetric reconstruction was performed using an adapted workflow in Scipion. We first reconstructed both proteins separately. The symmetric volume of the reference (Cep152) was then refined using the weighted sum of the input particle images (Cep152+2*HsSAS-6) without applying a symmetry constraint. Into the resulting asymmetric joint volume, we could fit the individual volumes to obtain a high-resolution dual color model. The volume fitting was performed using Chimera^30^.

### 2D particle averaging and volume alignment

Since the centriole has a principal rotational symmetry, we can collapse one of the spatial axes (x/y or z) to then identify the spatial translation parameters required to align two volumes. Specifically, we used particle projections of centriolar top (xy) or side (xz) views. In order to efficiently identify these specific orientations from a large number of individual particles, we calculated 12 parameters whose values could be used as a characteristic signature for top (xy) and side (xz) view projections. Next, a subset of 200 particles was selected and manually classified into top, side or intermediate views (i.e. the response). The shape descriptors and the manual classification were copied into a data table that can be used as a training dataset to generate models using supervised machine learning. We used MATLAB’s Classification Learner to identify the best model able to predict the classified outcome (response) based on the shape parameters. The best model was subsequently saved and could later be applied to other data sets. The model’s accuracy in predicting a certain shape worked in general better for top (xy) orientations, requiring little manual selection/filtering. Importantly, only one of the two imaged centriolar proteins (i.e. the reference) needs to be classified onto top/side view.

All of the following operations are then performed on both channel datasets. The identified particles were aligned to the center of mass of the reference protein and rendered using a pixel size of 10 nm. We next performed a rotational alignment using an extended version of efficient subpixel registration by cross-correlation^31^. Specifically, during the original procedure, we rotated each image stepwise from 1 – 359° by 1° at each iteration, resulting in 360 cross-correlations, from which we picked the orientation with the maximum root mean square (RMS) error, giving the optimal angle of rotation. The alignment was performed over ten iterations. The sum of all images was used as a reference for the first iteration. For all following iterations, we used the sum of all aligned particles from the previous iteration as the reference. The translation between both channels along the z axis was determined using a line profile measurement of the two-color reconstruction (Supplementary Fig. 9). To generate a final multi-color volume representation, the co-oriented volumes were loaded into Chimera^30^ and centered on top of each other. The z axial transformation was applied using the transform coordinates tool (Tools > Movement > Transform coordinates). The volume obtained from the lower resolution SMLM channel (i.e. DyLight755 channel, Supplementary Fig. 4) was then replaced by a higher resolution volume of the same structure. To this end, the high resolution volume was loaded into Chimera and aligned to the low resolution volume using the ‘Fit in Map’ tool (Tools > Volume Data > Fit in Map) and then transformed as described above.

### SMLM Simulations

In order to evaluate the contribution of labelling noise and efficiency as well as to test the particle processing workflow, we developed a particle simulator that generates localization maps from ground truth models. To define a starting structure, we generated ground truth models of expected fluorophore positions using geometric dimensions of the complex as obtained from SMLM. The starting structure was then randomly rotated and projected onto the XY plane. A random number of molecules were chosen according to the labeling efficiency and a defined number of noise molecules placed at random positions around each particle. Localizations (single frame appearance of a blinking event) originating from each fluorophore were simulated according to measured distributions for photon count, localization precision, as well as on- and off-time. The measurement distributions were obtained from single molecule calibrations for Alexa 647 performed under experimental conditions. The resulting simulated particles were analyzed as described for experimental SMLM data sets. All simulations were performed using custom-written Matlab code supplied as part of the supplementary software package.

### Code availability

All developed code is provided as Supplementary Software. Updates will be available from GitHub (https://github.com/christian-7/MultiColorSPR). Test data sets are available via Zenodo (https://doi.org/10.5281/zenodo.1127010).

## Acknowledgements

We thank Caroline Lehmann for growth and purification of T4 Bacteriophages, Jose Miguel de la Rosa Trevin for support regarding the optimal use of Scipion and Ciaran G. Morrison for providing the Cep164 antibody (1F3G10). N.B. was supported initially by a grant from the European Research Council (ERC) to P.G. (AdG 340227), and then by the EPFL Fellows postdoctoral fellowship program funded by the European Union’s Horizon 2020 Framework Programme for Research and Innovation (Grant agreement 665667, MSCA-COFUND). Research in S.M’s lab is supported by the National Centre for Competence in Research (NCCR) Chemical Biology and the ERC (243016-PALMassembly). K.M.D. is supported by a SystemsX.ch Transition Post-doc Fellowship (TPdF 2014/227). We thank Michal Daszykowski for providing a public DBSCAN implementation.

## Author Contributions

C.S, N.B, P.G, S.M. conceived and designed the project. C.S and S.M supervised the project. C.S. and N.B. performed all experiments and data analysis. C.S and K.M.D wrote analysis code. All authors wrote and revised the final manuscript.

## References

1. Campbell, M. G., Veesler, D., Cheng, A., Potter, C. S. & Carragher, B. 2.8 Å resolution reconstruction of the Thermoplasma acidophilum 20S proteasome using cryo-electron microscopy. Elife 4, e06380 (2015).

2. Jiang, J., Pentelute, B. L., Collier, R. J. & Zhou, Z. H. Atomic structure of anthrax protective antigen pore elucidates toxin translocation. Nature 521, 545–549 (2015).

3. Byeon, I.-J. L. et al. Structural convergence between Cryo-EM and NMR reveals intersubunit interactions critical for HIV-1 capsid function. Cell 139, 780–90 (2009).

4. Beck, M., Lučić, V., Förster, F., Baumeister, W. & Medalia, O. Snapshots of nuclear pore complexes in action captured by cryo-electron tomography. Nature 449, 611–615 (2007).

5. Strauss, M., Schotte, L., Karunatilaka, K. S., Filman, D. J. & Hogle, J. M. Cryo-electron Microscopy Structures of Expanded Poliovirus with VHHs Sample the Conformational Repertoire of the Expanded State. J. Virol. 91, e01443-16 (2017).

6. Chang, Y.-W. et al. Architecture of the type IVa pilus machine. Science 351, aad2001 (2016).

7. Szymborska, A. et al. Nuclear pore scaffold structure analyzed by super-resolution microscopy and particle averaging. Science 341, 655–8 (2013).

8. Salas, D. et al. Angular reconstitution-based 3D reconstructions of nanomolecular structures from superresolution light-microscopy images. Proc. Natl. Acad. Sci. 201704908 (2017). doi:10.1073/PNAS.1704908114

9. Douglass, K. M., Sieben, C., Archetti, A., Lambert, A. & Manley, S. Super-resolution imaging of multiple cells by optimized flat-field epi-illumination. Nat. Photonics 10, 705–708 (2016).

10. Bornens, M. The Centrosome in Cells and Organisms. Science (80-.). 335, 422–426 (2012).

11. Bauer, M., Cubizolles, F., Schmidt, A. & Nigg, E. a. Quantitative analysis of human centrosome architecture by targeted proteomics and fluorescence imaging. EMBO 35, 1–15 (2016).

12. Kitagawa, D. et al. Structural Basis of the 9-Fold Symmetry of Centrioles. Cell 144, 364–375 (2011).

13. Gönczy, P. Towards a molecular architecture of centriole assembly. Nat. Rev. Mol. Cell Biol. 13, 425–35 (2012).

14. Ester, M., Ester, M., Kriegel, H.-P., Sander, J. & Xu, X. A density-based algorithm for discovering clusters in large spatial databases with noise. Proc. 2nd Internat. Conf. Knowl. Discov. Data Min. 226-231 (1996).

15. Rosa-Trevin, de la, J. M. et al. Scipion: A software framework toward integration, reproducibility and validation in 3D electron microscopy. J. Struct. Biol. 195, 93–99 (2016).

16. Scheres, S. H. W. et al. Maximum-likelihood Multi-reference Refinement for Electron Microscopy Images. J. Mol. Biol. 348, 139–149 (2005).

17. Lukinavičius, G. et al. Selective chemical crosslinking reveals a Cep57-Cep63-Cep152 centrosomal complex. Curr. Biol. 23, 265–270 (2013).

18. Momotani, K., Khromov, A. S., Miyake, T., Stukenberg, P. T. & Somlyo, A. V. Cep57, a multidomain protein with unique microtubule and centrosomal localization domains. Biochem. J. 412, 265–73 (2008).

19. Sonnen, K. F., Schermelleh, L., Leonhardt, H. & Nigg, E. a. 3D-structured illumination microscopy provides novel insight into architecture of human centrosomes. Biol. Open 1, 965–76 (2012).

20. Banterle, N. & Gönczy, P. Centriole Biogenesis: From Identifying the Characters to Understanding the Plot. Annu. Rev. Cell Dev. Biol. 33, 23–49 (2017).

21. Loncarek, J., Hergert, P., Magidson, V. & Khodjakov, A. Control of daughter centriole formation by the pericentriolar material. Nat. Cell Biol. 10, 322–328 (2008).

22. Gogendeau, D., Guichard, P. & Tassin, A.-M. in Methods in cell biology 129, 171–189 (2015).

23. Bornens, M., Paintrand, M., Berges, J., Marty, M.-C. & Karsenti, E. Structural and chemical characterization of isolated centrosomes. Cell Motil. Cytoskeleton 8, 238–249 (1987).

24. Bourdin, G. et al. Amplification and purification of T4-like escherichia coli phages for phage therapy: from laboratory to pilot scale. Appl. Environ. Microbiol. 80, 1469–76 (2014).

25. Edelstein, A. et al. in Current Protocols in Molecular Biology 14.20.1-14.20.17 (John Wiley & Sons, Inc., 2010). doi:10.1002/0471142727.mb1420s92

26. Olivier, N., Keller, D., Gönczy, P. & Manley, S. Resolution doubling in 3D-STORM imaging through improved buffers. PLoS One 8, e69004 (2013).

27. Huang, F. et al. Video-rate nanoscopy using sCMOS camera–specific single-molecule localization algorithms. Nat. Methods 10, 653–658 (2013).

28. Douglass, K. M., Sieben, C., Berliner, N. & Manley, S. B-Store. (2017). doi:10.5281/ZENODO.1117843

29. Nieuwenhuizen, R. P. J. et al. Measuring image resolution in optical nanoscopy. Nat. Methods 10, 557–62 (2013).

30. Pettersen, E. F. et al. UCSF Chimera – A visualization system for exploratory research and analysis. J. Comput. Chem. 25, 1605–1612 (2004).

31. Guizar-Sicairos, M., Thurman, S. T. & Fienup, J. R. Efficient subpixel image registration algorithms. Opt. Lett. 33, 156 (2008).

